# Removal modelling in ecology

**DOI:** 10.1101/2020.02.20.957357

**Authors:** Oscar Rodriguez de Rivera, Rachel McCrea

**Affiliations:** Statistical Ecology @ Kent, National Centre for Statistical Ecology. School of Mathematics, Statistics and Actuarial Science, University of Kent, Canterbury, CT2 7FS, UK

## Abstract

Removal models were proposed over 80 years ago as a tool to estimate unknown population size. Although the models have evolved over time, in essence, the protocol for data collection has remained similar: at each sampling occasion attempts are made to capture and remove individuals from the study area. Within this paper we review the literature of removal modelling and highlight the methodological developments for the analysis of removal data, in order to provide a unified resource for ecologists wishing to implement these approaches. Models for removal data have developed to better accommodate important feature of the data and we discuss the shift in the required assumption for the implementation of the models. The relative simplicity of this type of data and associated models mean that the method remains attractive and we discuss the potential future role of this technique.

**Author summary:** Since the introduction of the removal in 1939, the method has being extensively used by ecologists to estimate population size. Although the models have evolved over time, in essence, the protocol for data collection has remained similar: at each sampling occasion attempts are made to capture and remove individuals from the study area. Here, we introduce the method and how it has been applied and how it has evolved over time. Our study provides a literature review of the methods and applications followed by a review of available software. We conclude with a discussion about the opportunities of this model in the future.

## Introduction

One of the most interesting problems for the ecologist is the estimation of the density of a particular animal species [1] and a tremendous diversity of techniques are available for estimating the size of populations [2–4]. Despite frequent refinements to make some of the more sophisticated techniques more realistic for particular situations (see for example, [5–9]) the removal method remains exceptionally popular.

Removal (or depletion) sampling is a commonly used method to estimate abundance of animal populations [10–13]. Removal models have been used to estimate population size, not only for many species including birds [14], mammals [15], and fish [16], but also for epidemiological applications [17–19].

Removal models are ideally suited to estimating the number of invasive species as they coincide with desirable management (i.e. the reduction or eradication of populations) [20] and the method has recently been adopted as a conservation management tool for example for mitigation translocations [12, 21]. Models that use data from management actions need to account for variations in removal effort as these data are unlikely to be standardised across events [20]. [22] showed that removal models that account for removal effort are effective at estimating abundance, particularly when removal rates are high.

The classic removal model was introduced by [23] and [24], motivated by a theory developed by [1]. This model relied on the assumption of population closure and constant detection probability, meaning that the animals are assumed to be available for capture with the same probability throughout the study and there are no births, deaths or migration during the study. The basic removal model results in a geometric decline in the expected number of captured individuals over time. This classic removal model is a special case of model *M*_*b*_ for closed populations which allows for a behavioural response to initial trapping [5].

### Overview of the paper

Within this paper we have conducted a systematic literature review of removal modelling in ecology. We describe the methods applied in the systematic review and the aspects of interest. We present the results obtained from the literature analyses, highlighting the key methodological advances which have been made in this field and a review of software which has been used to fit removal models. The paper concludes with a discussion about the future role of removal modelling in ecology.

## Materials and methods

### Literature Search

This systematic review followed the PRISMA (Preferred Reporting Items for Systematic Reviews and Meta-Analyses), statement as a guide [25]. The bibliographic search was performed using the SciVerse Scopus (https://scopus.com), ISI Web of Science (https://webofknowledge.com), and Google Scholar (https://scholar.google.com) databases. Papers published between 1939 and the cut-off date 01 July 2019 with the terms “Removal model” or “Removal method” and “population” in the title, keywords, or abstract were included. Non-English publications, and papers reporting removal methods focused on cleaning procedures were excluded from our search. The process of selecting papers to include in our review started with a screening of the abstract. Articles were excluded if they mentioned the keyword “removal model/method” for justification or discussion without implementing a removal model as part of the study. Thus, only the papers that reported applications, methodological advances of removal models or removal study design were retained for the analysis (Fig 1).

**Fig 1.**
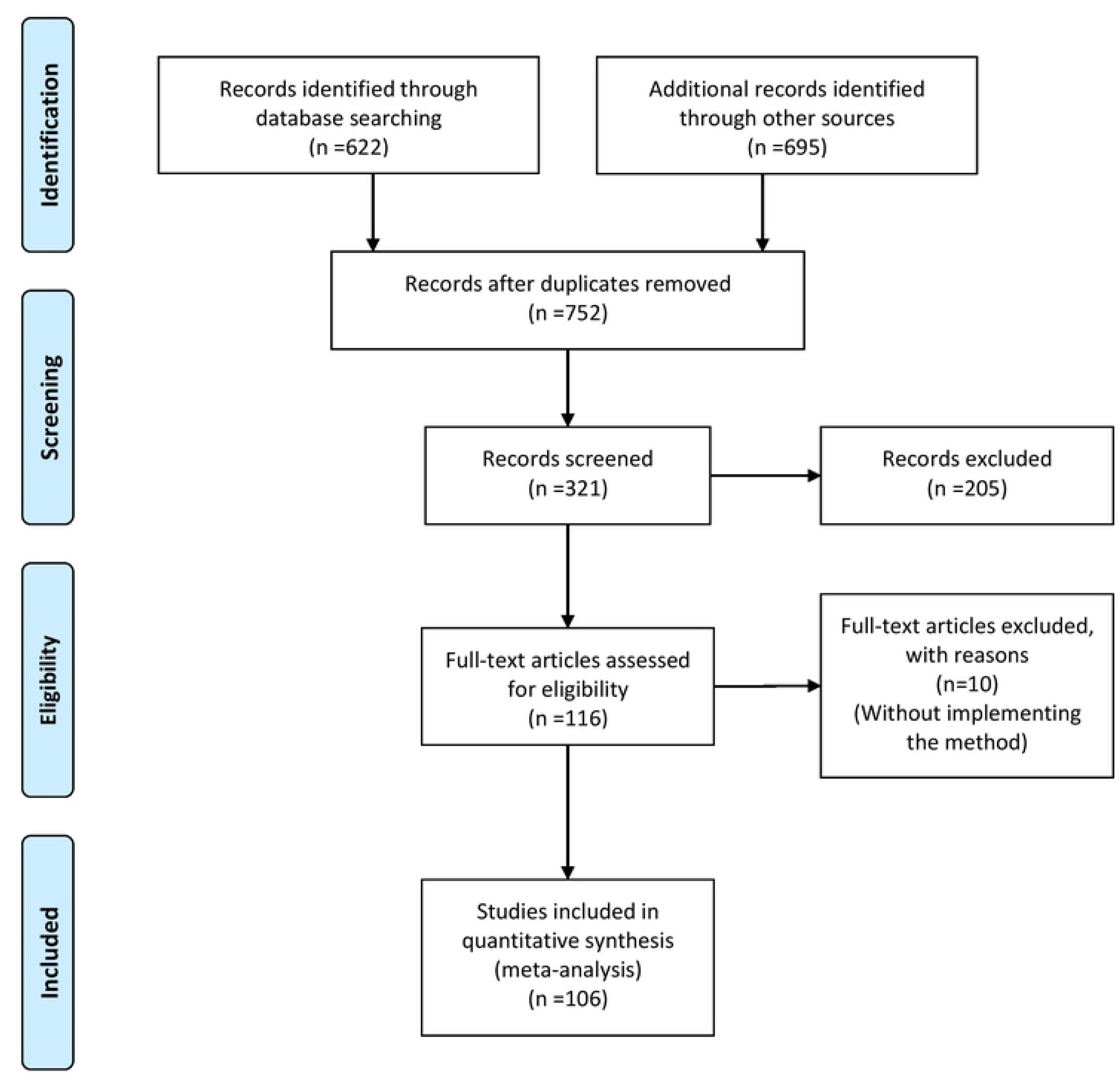
PRISMA flowchart.

### Data Analysis

Features and parameters of each study were categorised and compiled into a database of removal model publications. We identified the details that were reported in the 104 reviewed publications and provided a descriptive summary of the essential details that need to be reported in published removal models (see Table 1).

**Table 1.**
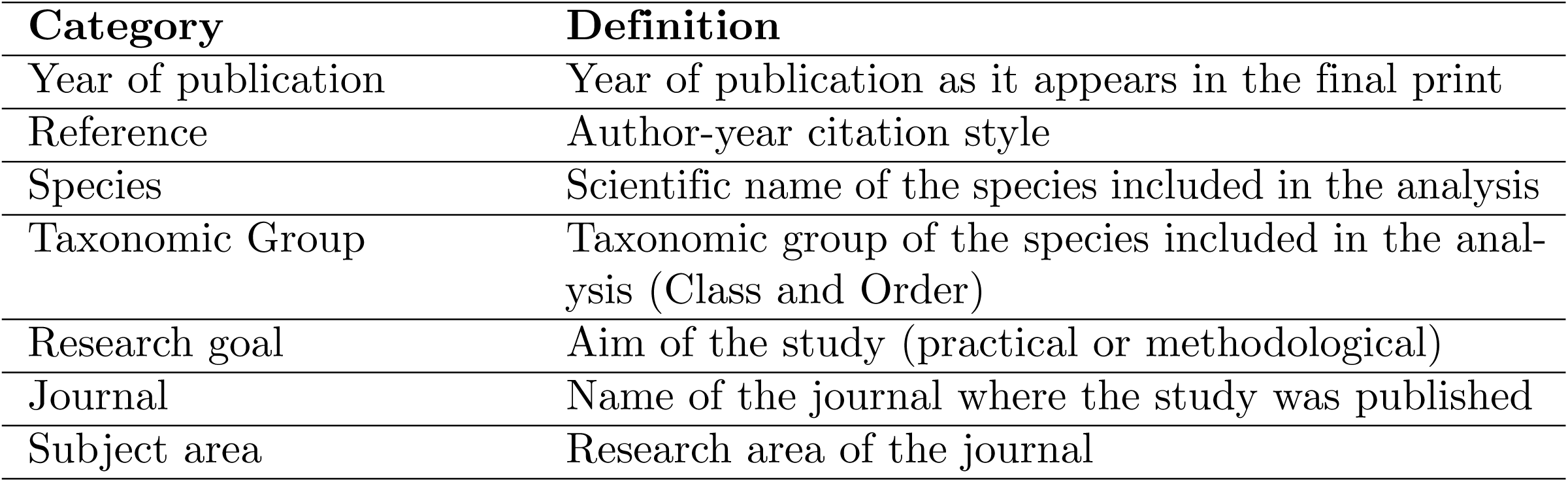
Parameters used to categorise Removal models included within the database

## Results

### Synthesised Findings

The reviewed literature was published from 1939 to 2019 and interestingly there have historically been long gaps in publications on this topic. However in recent years there has been a more constant stream of published papers, suggesting a resurgence of interest in the method (Fig 2). This is potentially an indication of the role of removal modelling in studies of reintroduction, especially when translocated individuals are removed from endangered populations [26], and the adaptations of model collection protocols to adopt a “removal” design for other data types such as occupancy and distance sampling, as will be discussed in “Adapting sampling schemes using removal theory” Section.

**Fig 2.**
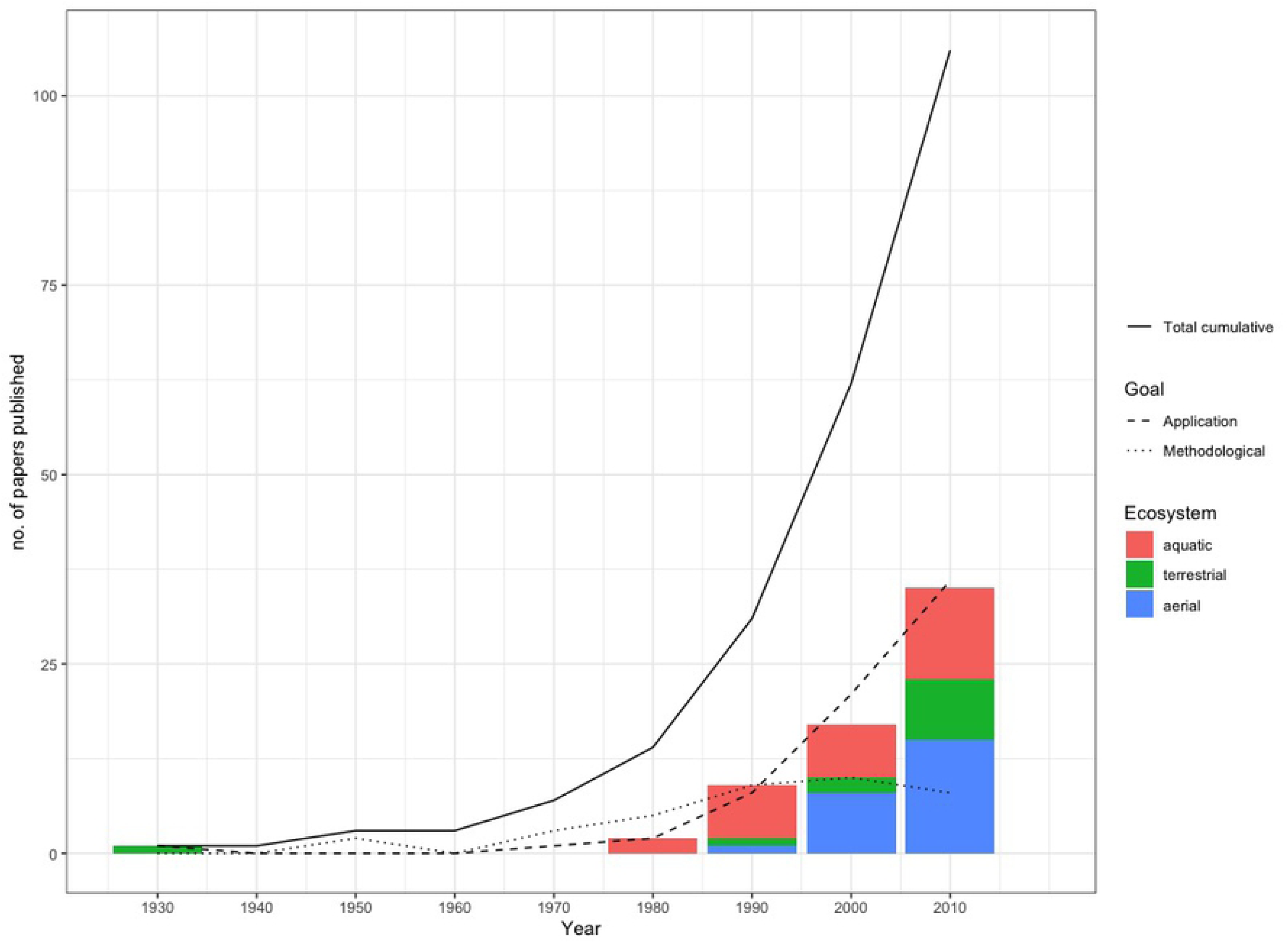
Papers published since 1939 by category. Papers published since 1939 by category (application or methodological) and ecosystem (aquatic, terrestrial or aerial

A strong representation of removal model publications was observed in the fields of Statistics (29 publications), Fisheries and Ecology journals (20 papers each of the disciplines) and 13 papers published in ornithological journals.

### Methodological Contributions

#### Early model developments

The basic principle of the removal method is that a constant sampling effort will remove a constant proportion of the population present at the time of sampling. Thus, if the total population size is *N* and *p* denotes the probability of capture of an individual, the expected number of captures will be given by: *pN, p*(*N* − *pN*) and *p*[*N* − *pN* − *p*(*N* − *pN*)] for the first, second and third sampling occasions, respectively. Population estimates can be obtained either by plotting catch per unit effort of collection as a function of total previous catch (see for example [1, 27, 28]) or by obtaining maximum likelihood estimates [23, 24, 29, 30].

This model can be formalised by defining the corresponding likelihood function. Suppose *x*_*t*_ denotes the number of individuals captured at sampling occasion *t* = 1, …, *T*, where *T* denotes the number of sampling occasions.

Using a binomial-formulation, if *N*_*t*_ individuals are still in the study at sampling occasion *t*, we can define the probability of removing *x*_*t*_ individuals by

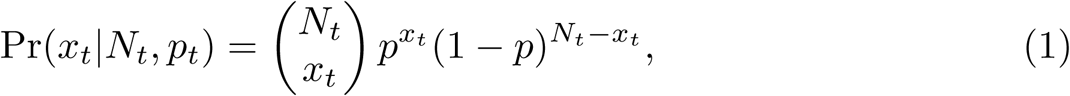

where *N*_1_ = *N* and 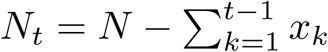 for *t* ≥ 2. Then, the probability of observing *x*_1_, …, *x*_*T*_ is given by

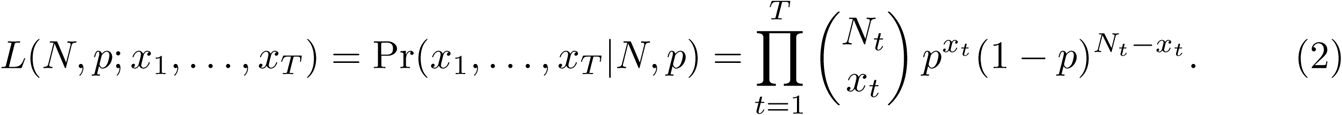

Alternatively, we could specify that the *N* individuals within the population belong to one of *T* + 1 categories: either they are captured on one of occasion 1, …, *T*, or they could never be captured. Let *π*_*t*_ denote the probability that an individual is captured at occasion *t*

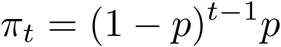

and let *π*_0_ denote the probability that an individual is never captured:

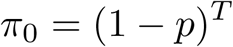

Let *n* denote the number of individuals never captured, which is given by 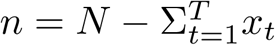, then the likelihood can be expressed as:

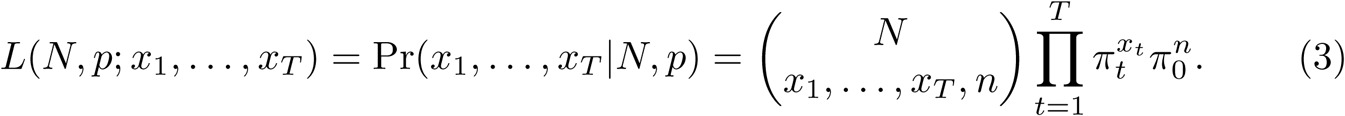

Early developments of removal models were methodological in nature, to overcome issues which these days are simple to deal with because of computing power. [24] formalised the conditional binomial removal model of [23], providing an asymptotic variance of the abundance estimator. Further, they demonstrated how graphical methods can be used to estimate the parameters of capture and abundance. The likelihood of [23] is weighted with a beta prior by [31], which results in estimators with lower bias and variance. [32] proposed an improved confidence interval for abundance for small populations, whilst [33] proved that the profile log-likelihood for the removal model is unimodal and demonstrated that the likelihood-ratio confidence interval for the population size has acceptable small-sample coverage properties. Similarly, [34] proposes a profile likelihood approach for estimating confidence intervals which showed improved performance.

#### Validity of model assumptions

The model assumptions required for these early models were very restrictive. Specifically, capture probability, *p* is assumed to be constant both across all individuals and for each sampling period. [23, 24, 29, 30] have typically included tests of assumptions or information on necessary sample sizes in their studies. [13] included a table of percentage errors to be expected if *p* varies during sampling and [5] offered alternative ways to estimate *N* if *p* varied for a wide class of closed population capture-recapture models, however it is not possible to fit many of these to removal data which has only one occasion of capture. [35], noting that the assumptions of the removal model are often violated, proposed the non-parametric jackknife estimator as an alternative to the removal model.

[31] proposed a standard test that combines testing for addition or deletions to the population with testing for equal catchability. The test entails examining trends in the catch vectors: “When the expected third catch as determined from the first two catches is larger than the observed third catch, emigration or a decreasing probability of capture is indicated. When the opposite condition exists, immigration or an increase in the probability of capture is indicated”. Both the presence and absence of trends yields ambiguous information. Significant trends in the catch vectors can be caused by the population being open or by unequal catchability. An absence of trends implies either that the assumptions were met or that migration balanced changing probabilities of capture over time.

[36] indicates that unequal catchability tends to be the rule in biological populations. Hence testing for equal catchability is crucial unless one adapts the model. Assuming that equal catchability exists when it does not leads to underestimation of the population size. One procedure that avoids this bias is to identify subsets of the population that are equally catchable and to obtain separate population estimates for each subset [37]. However, because such subdivision of the data greatly decreases the precision of the estimate of the total population size, one should not divide the population unnecessarily if the assumption of equal catchability of all the individuals is met. Therefore, employing a test of equal catchability is a crucial step in any population analysis, even if failure to reject the null hypothesis of equal catchability is an ambiguous result [38].

Two aspects of equal catchability are important for the removal method: equal catchability among groups and equal catchability in all sampling occasions. The first is tested analogously to the test for marked and unmarked captures. If groups with different catchability are identified, separate population size estimates are made for each such group. Individual differences in catchability unrelated to a particular group membership are still possible, but [32] showed that unless these differences are great, their effect on the population estimate is small. [39] investigated the robustness of the removal model to varying behaviours exhibited by fish using simulation.

The second assumption, that catchability remains constant in all sampling periods can be tested by the chi-square test given by [24] or further test given by [5] or [40]. Conclusions drawn from any of these tests will be accurate only if the population remains closed during sampling. Use of a barrier if possible, is again desirable, or independent verification by sampling of marked animals [37].

The closure assumption of the removal model has been relaxed in [41], where a model was proposed which allows for population renewal through birth/immigration as well as for population depletion through death/emigration in addition to the removal process. The arrivals of new individuals are modelled by an unknown number of renewal groups and a reversible jump MCMC approach is used to determine the unknown number of groups. Within this paper however it is assumed that any emigration from the population is permanent, and this assumption has been relaxed in [12] which presents a robust design, multilevel structure for removal data using maximum likelihood inference. The implemented hidden Markov model framework [42], allows individuals to enter and leave the population between secondary samples. A Bayesian counterpart to this robust design model is presented in [43].

#### Change in ratio, index-removal and catch-effort models

Change in ratio models for population size estimation are closely related to removal models [44, 45]. The model requires that the population can be sub-divided into distinct population classes and the removals will be performed in such a way that the ratio of removals of the sub-populations will be the same as the underlying ratio of the sub-classes within the population.

We can generalise the basic removal model likelihood function of Eq (2) by extending the definition of the parameters and the summary statistics. Specifically, suppose the population is sub-divided into *G* mutually exclusive and exhaustive groups, and let *N* (*g*) denote the unknown abundance of sub-population *g* = 1, …, *G*. We now record *x*_*t*_(*g*) individuals of sub-population *g* being removed at occasion *t*. The likelihood becomes,

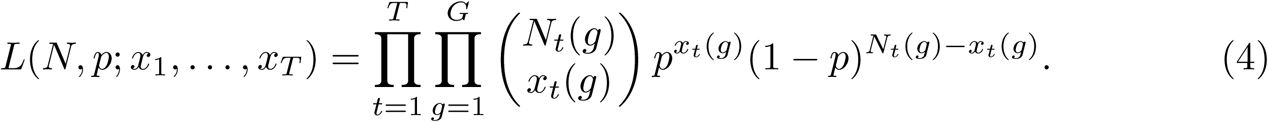

where *N*_1_(*g*) = *N* (*g*) and 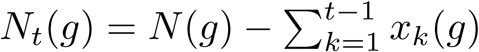 for *t* ≥ 2.

Catch-effort models are a straight-forward extension of basic removal models which allow capture probability to be related to sampling effort. If catch per unit effort declines with time, then regressing accumulated removals by catch per unit effort allows the starting population to be estimated. This approach however strongly relies on the assumption that if more effort is put into capturing the individuals then a higher proportion of the population will be caught and if this is not satisfied estimators might be appreciably biased [46]. More generally we can extend the removal likelihood of Equation 2 to define a functional form of capture probability:

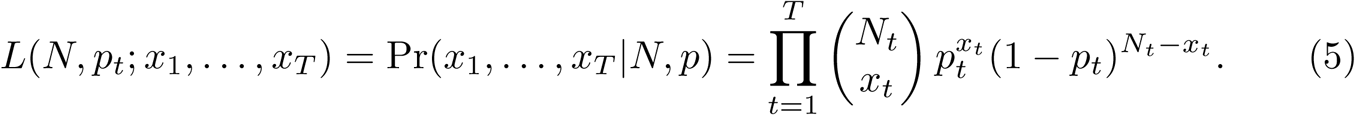

where *p*_*t*_ is the capture probability at occasion *t* which can be linked to a recorded covariate of survey effort, denoted by *w*_*t*_. Possible forms for the functional form might be logit(*p*_*t*_) = *α* + *βw*_*t*_, where *α* and *β* are parameters to be estimated, or 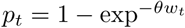, where *θ* is a single parameter to be estimated has been used for fisheries applications where *w*_*t*_ denotes the amount of time spent fishing. Alternatively if *w*_*t*_ denotes the number of traps on occasion *t* and each animal is assumed to be caught in any trap with probability *θ*, 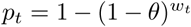 [4]. Indeed the logistic form of time-dependent capture probability can also be used to model time-variation in capture probability as a function of climatic conditions - see for example [47].

When sampling is with replacement and the sampling efforts are known, [48] modelled the survey sampling process as a Poisson point process where each animal is counted at random with respect to increments of sampling effort and it is assumed that the encounter probabilities for each individual are independent. [49] propose a class of catch-effort models which allow for heterogeneous capture probabilities.

The index-removal method makes use of the decline in a measure of relative abundance due to a known removal. The relative abundance is measured in surveys before and after the removal [50]. [51] proposed an index-removal estimator which accounts for seasonal variation in detection.

#### Further model developments

[52] demonstrates why auxiliary information is beneficial in removal studies and how to incorporate the extra information into the model and [14] extended the idea of incorporating sub-class level information by proposing a conditional likelihood approach for incorporating auxiliary variables. Capture probabilities are directly estimated from the conditional likelihood and then abundance estimates can be obtained using a Horvitz-Thompson-like approach.

[53] relaxed the assumption of [23] that traps are not limited in capacity by developing a model in which traps have a reduced capacity to catch once they have been filled. The model assumes that the probability that a specific animal will be caught is proportional to the number of traps that are unoccupied and is often referred to as the proportional trapping model.

Continuous-time removal models were proposed in [54] and the proportional trapping model and continuous time framework were combined in [55]. Further [56] and [57] extended the continuous time proportional trapping model to account for known ratios of sub-populations, thus generalising the change-in-ratio approach.

The theory of analysing multiple types of data in an integrated model within ecology has gained traction in recent years - see for example [58]. Early ideas of combining data types has been found in the removal literature. For example, [37] proposed combining capture-recapture and removal methods for fish removals when sampling is over a limited study period and [59] showed how the proportional trapping model can be extended to include data on non-target species.

Removal models have been presented as a class of hierarchical models, for example [16] present a hierarchical removal model where the sites are assumed to have several distinct sub-sites located spatially. Suppose removals occur at *S* sites, then records are made of *x*_*st*_, the number of individuals removed from site *s* = 1, …, *S* at occasion *t* = 1, …, *T*. Following the multinomial form of the likelihood of Equation 3 the probability of observing a sequence of removal counts, **x**_**s**_ = {*x*_*s*1_, …, *x*_*sT*_} from site *s* is given by

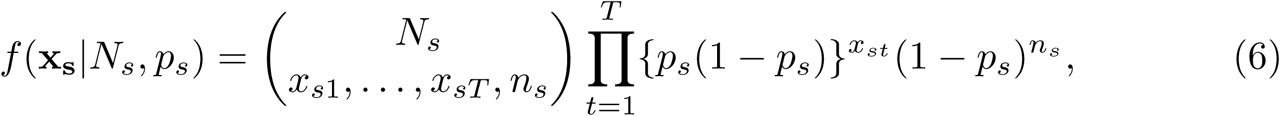

where *N*_*s*_ denotes the abundance at site *s, p*_*s*_ denotes the capture probability at site *s* and 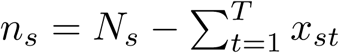. Within the hierarchical formulation, a probabilistic formulation, defined by density function *g*(*N*|*ψ*), with parameter *ψ*, specifies the variation in abundance among the *S* spatially distinct sub-populations in the sample. The site-specific removal counts (Equation 6) can be combined with this model by integrating over possible values of *N*_*s*_:

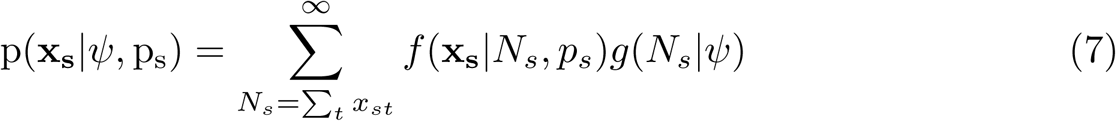

The likelihood function is then the product over the observations from the *S* sites, which assuming independence is defined by

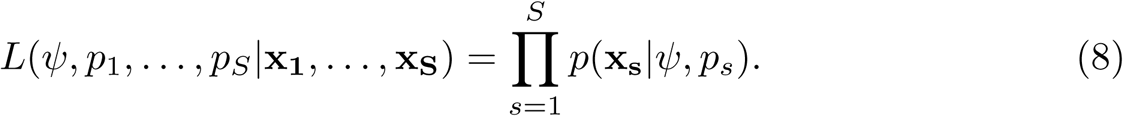

This model is in fact a multinomial N-mixture model [60] and it has been shown that the removal N-mixture model outperforms the standard N-mixture model using simulation [61]. In practice, a value *K* has been used in place of the infinite sum in equation 8 when evaluating the likelihood, however what value of *K* is appropriate is subjective, and it has been shown there can be some problems with proposed values [62]. [63] has demonstrated that the multinomial N-mixture model for removal data, with negative binomial mixing distribution, has a closed-form likelihood and therefore no numerical approximations are required to fit the model.

#### Adapting sampling schemes using removal theory

Little work has been found which investigates study design for removal surveys. [64] explored how to optimally allocate total sampling effort for multiple removal sites by maximising the Fisher information of the constant capture probability in the classic removal model. This approach was extended by [65] to allocate effort between primary and secondary sampling occasions within the robust design removal model [12].

An adaptation of survey design for various data types have been augmented with the concept of removal studies. For example, [66] described a time-removal model that treats subsets of the survey period as independent replicates in which birds are ‘captured’ and mentally removed from the population during later sub-periods. This method has been implemented in many subsequent papers, see for example [67, 68]. Further, the removal study design has been proposed for occupancy surveys [69], whereby once a site has been observed as occupied by a species no further surveys are required [70]. [71] developed a spatially explicit temporary emigration model permitting the estimation of population density for point count data such as removal sampling, double-observer sampling, and distance sampling.

### Applications

From the papers we reviewed three ecosystems were identified (Figure 2) based on the species analysed. Almost half of the applied studies were focused on aquatic ecosystems (marine and fresh water); followed by flying species (n=24) and the rest of the applied studies analysing terrestrial ecosystems. However partitioning the papers by taxonomic group shows that the most common group are bird applications, with 23 papers identified. However, it should be noted that many of these applications use other data types but with adapted sampling design as described earlier [14, 66, 72–92]. Removal methods are clearly important in fisheries research and applications are presented in [93–110]. The papers analysing data from mammals are [1, 5, 20, 111–121]. There are a further six papers analysing data from amphibians: [122–127] and [51] analysed crustaceans. Insects have being analysed in three papers: [128–130] and the less common applications included annelids [131] and Holothuroidea [132]. Three papers presented applications about human disease [17–19] and we included these in our analysis as the aim of the study was to estimate the proportion of an affected population which is an aim in common with ecological applications.

### Software

There has been a considerable amount of recent work on developing software to make complex statistical models accessible to the wider ecological community. Much software has been developed to estimate population parameters, including abundance and demographic parameters which account for imperfect detection.

`Capture` [5], was developed to compute estimates of capture probability and population size for closed population capture-recapture data and given the basic removal model is a special case of closed capture-recapture model *M*_*b*_, `Capture` can be used to fit the geometric removal model. `RCapture` [133] is an R package [134] for capture-recapture experiments. `Rcapture` can fit models for capture-recapture models but as well as standard open, closed and robust-design versions of these models based on multinomial likelihoods it is the only software which also implements a log-linear modelling algorithm to estimate the parameters.

`Program Mark` [135] provides a wide-range of models which can be fitted to more than 65 different data types to estimate several population parameters from the encounters of marked animals. Typically, parameters are obtained by method of maximum likelihood estimation through numerical methods (Newton-Raphson by default). However, an MCMC algorithm has been added to provide estimates using a Bayesian framework [136]. [137] demonstrate how removal models can be fitted using *Program Mark* for fisheries data. `RMark` [138] is a software package for the R computing environment that was designed as an alternative interface that can be used in place of `Program Mark’s` graphical-user-interface to describe models with a typed formula so that models do not need to be defined manually through the design matrix. At the time of writing `RMark` supports fitting 97 of the 155 models available in *Program Mark*. R packages `marked` [139] and `unmarked` [140] can also be used to fit the standard removal model and multinomial N-mixture removal model, respectively - see [61]. There is also more specialised software that has arisen for specific applications. `Removal Sampling` v2 [141] was designed to estimate population size from removal data and [105] apply this software in order to analyse the effectiveness of stream sampling methods for capturing invasive crayfish.

In addition there is of course well-known software which accommodates the removal study design when fitting models to other data types. In particular `Presence` and `RPresence` [142] for occupancy surveys and `Distance` [143] for distance sampling.

## Discussion

Early removal models were simplistic and did not adequately account for potential variability exhibited by the underlying population, however computational and methodological advances give the possibility of more complex models, increasing opportunities, scenarios and accuracy in population estimation. The framework we have presented here is designed to assimilate the use of removal models in order to assist future practitioners in the effective application of removal models.

In our review we have not only presented a list of papers published, differentiating application and methodological advances, but also we have explained the evolution of the model. We have shown how the model has been developed since [1] presented the first case with the evolution in likelihood function from the basic to, for example, that proposed by [49], accounting for heterogeneous capture probability, and more recently the work of [63], theoretically developing the multinomial N-mixture model for removal data.

We have shown how the model assumptions have been adapted, trying to fit the model to different scenarios such as unequal catchability [36], non closure of population [41], heterogenity accross sites [16] and temporary migration [12, 43]).

Software development, means that even the complex models described in this paper are accessible to ecologist, meaning that maximum utility can be obtained from removal data.

There are several advantages for non-specialists that wish to apply removal methods: there is a vast array of available models for removal data, with the possibility of selecting the approach where the model assumptions best align with their particular study; there is no restriction to frequentist or Bayesian paradigms; there are several software packages and R code accompanying publications of more recent development to investigate where model assumption might fall short.

In our research we noted in earlier papers a thorough assessment of effects if model assumptions were violated but this rigour was not found in late methodological papers, except in some cases through part of a simulation. New methods developed in this research field have been motivated by unique aspects of particular data sets, and therefore nuances of a case study should be embraced rather than avoided in order to encourage methodological advances.

There is a worldwide interest in identifying tools for effective estimation of species population size and removal models show great potential for application in a wide range of situations, such as species relocation projects.

The potential of removal models to facilitate the estimation of population size in the source population whilst also obtaining a pool of individuals to translocate/reintroduce means that such models will remain important and will likely be further developed.

Species relocations are becoming more prevalent in conservation worldwide [144–146]. They are performed in several countries on an extensive range of species including plants [146], amphibians and reptiles [147, 148]. There are many studies of translocated species and the success of reintroductions, including settlement, survival and reproduction of translocated individuals and their effects on the viability of the reintroduced population [149–154]. However, there is less information regarding the impact of translocations on the source or donor population [155, 156]. These impacts can dramatically affect community stability, which is especially important when translocated individuals are from endangered populations [26]. The main components that can affect the stability of a population are: resistance, that is the ability to maintain its current state when subjected to a perturbation [157]; amplitude, that will determine, after some alteration, if it will return to its original state [158]; elasticity is the property that will determine the rate of return to its initial configuration when the perturbation exceeds the resistance of a community, but not its amplitude [158]. Removal data and removal models may be a powerful tool in order to understand and manage these populations.

